# Profiling and leveraging relatedness in a precision medicine cohort of 92,455 exomes

**DOI:** 10.1101/197889

**Authors:** Jeffrey Staples, Evan K. Maxwell, Nehal Gosalia, Claudia Gonzaga-Jauregui, Christopher Snyder, Alicia Hawes, John Penn, Ricardo Ulloa, Xiadong Bai, Alexander E. Lopez, Cristopher V. Van Hout, Colm O’Dushlaine, Tanya M. Teslovich, Shane E. McCarthy, Suganthi Balasubramanian, H. Lester Kirchner, Joseph B. Leader, Michael F. Murray, David H. Ledbetter, Alan R. Shuldiner, George Yancoupolos, Frederick E. Dewey, David J. Carey, John D. Overton, Aris Baras, Lukas Habegger, Jeffrey G. Reid

## Abstract

Large-scale human genetics studies are ascertaining increasing proportions of populations as they continue growing in both number and scale. As a result, the amount of cryptic relatedness within these study cohorts is growing rapidly and has significant implications on downstream analyses. We demonstrate this growth empirically among the first 92,455 exomes from the DiscovEHR cohort and, via a custom simulation framework we developed called SimProgeny, show that these measures are in-line with expectations given the underlying population and ascertainment approach. For example, we identified ∼66,000 close (first- and second-degree) relationships within DiscovEHR involving 55.6% of study participants. Our simulation results project that >70% of the cohort will be involved in these close relationships as DiscovEHR scales to 250,000 recruited individuals. We reconstructed 12,574 pedigrees using these relationships (including 2,192 nuclear families) and leveraged them for multiple applications. The pedigrees substantially improved the phasing accuracy of 20,947 rare, deleterious compound heterozygous mutations. Reconstructed nuclear families were critical for identifying 3,415 *de novo* mutations in ∼1,783 genes. Finally, we demonstrate the segregation of known and suspected disease-causing mutations through reconstructed pedigrees, including a tandem duplication in *LDLR* causing familial hypercholesterolemia. In summary, this work highlights the prevalence of cryptic relatedness expected among large healthcare population genomic studies and demonstrates several analyses that are uniquely enabled by large amounts of cryptic relatedness.

## Introduction/Background

The number and scale of large human sequencing projects is rapidly growing, including DiscovEHR^1^, UK Biobank^2^, the US government’s All of Us (part of the Precision Medicine Initiative)^3^, TOPMed (Web Resources), ExAC/gnomAD^4^, and many others. Many of these studies are collecting samples from integrated healthcare populations that have accompanying phenotype-rich electronic health records (EHRs) with a goal of combining the EHRs and genomic sequence data to catalyze translational discoveries and precision medicine^1^. These large-scale healthcare population-based genomic (HPG) studies are recruiting participants through healthcare systems where volunteers donate DNA and provide medically relevant metrics recorded in their EHRs. A major difference between the new HPG study design and traditional population-based studies is ascertainment, both in how participants are recruited and in the proportion of the population within in a geographical area that participates (Figure 1A). Traditionally, the high-expense of large-scale genetic studies and the limited resources of individual investigators has generated study populations exhibiting shallow ascertainment of individuals from a variety of geographical areas. To improve statistical power, samples from many different collection centers are combined into a larger cohort, and these cohorts are often merged into a much larger consortium consisting of tens to hundreds of thousands of individuals. While the total number of individuals sampled is often high, these studies typically only sample a relatively small portion of individuals in any given geographic area. In contrast, planned and on-going HPG studies are sampling tens to hundreds of thousands of participants from individual healthcare systems^1^.

**Figure 1.**
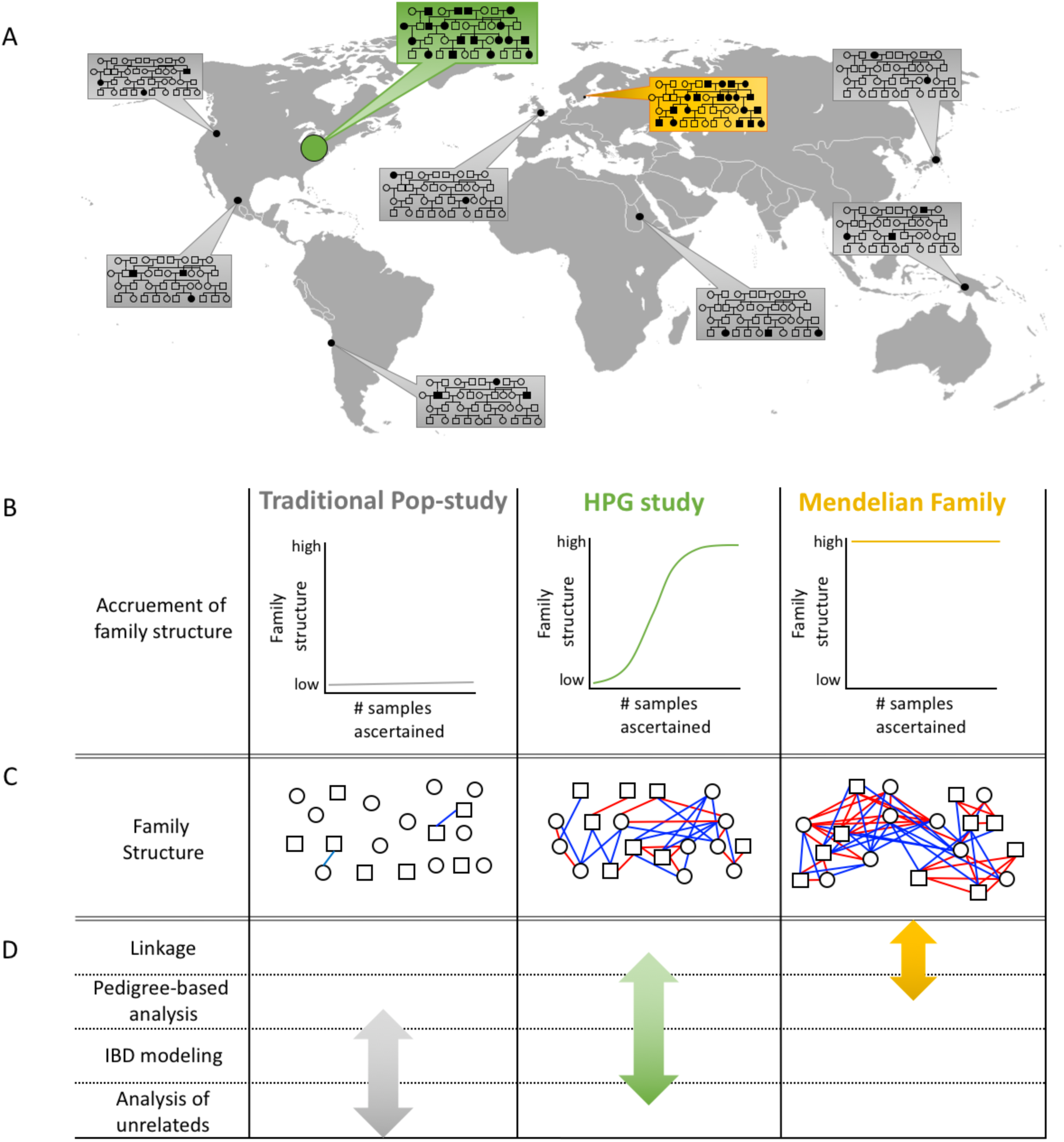
*Ascertaining a high proportion of the population in a geographical area increases family structure and impacts statistical analysis approaches that should be used. (A) Traditional population-based studies (gray boxes) typically sample a small portion of individuals from several populations. HPG studies (green box) more densely sample individuals from one or more populations. Family-based studies (yellow box) heavily sample within extended families, but do not sample nearly as many individuals as the other two study designs. (B) The three study designs result in very different levels of individuals with one or more close relatives in the dataset. (C) These three ascertainment approaches result in very different amounts of family structure. Red and blue lines indicate first- and second-degree pairwise relationships, respectively. HPG studies are expected to contain a level of family structure between the other two designs. (D) For this study, statistical analysis approaches were binned into four categories based on the level of family structure required to effectively use the approach (first column): “Linkage” refers to traditional linkage analyses using one or more informative pedigrees; “Pedigree-based analysis” refers to statistical methods beyond linkage that use pedigree structures within a larger cohort that includes unrelated individuals; “IBD modeling” refers to analysis that model the pairwise relationships between individuals without using the entire pedigree structure; “Analysis of Unrelateds” refers to analyses that assume all individuals in the cohort are unrelated. The amount of family structure impacts the approaches that can be used, and the arrows indicate the analysis ranges for which the three study designs are best suited.*

The difference in these two ascertainment approaches results in different patterns of genetic relatedness among individuals in these cohorts. Relatedness is a continuum that manifests itself within a cohort in a variety of ways depending on the population and how individuals are sampled from it. Because traditional population-based studies have generally collected samples from multiple geographical areas, they most commonly exhibit the broadest “class” of relatedness: *population structure*. Population structure (often referred to as “substructure” or “stratification”) within a genetic study results from it containing different ancestral groups or “genetic demes” in which allele frequencies are more similar within genetic demes than between demes. Genetic demes arise due to more recent genetic isolation, drift, and migration patterns. Classic examples of homogenous genetic demes include founder populations such as the Ashkenazi Jewish^5,6^ and Old Order Amish^7^, but they also occur at many levels of genetic isolation such as continental, sub-continental^6,8,9^, and even within the same urban community^5,10^. When genetic demes begin to mix, they create genetically admixed populations. Studies that contain an admixed population or more than one genetic deme are likely to contain population structure. Ascertainment of individuals within genetic demes can generate *distant cryptic relatedness*^5,6^, the second “class” of relatedness^11^, defined here as third-to ninth-degree relatives. These distant relatives are unlikely to be identifiable from the EHR but are important because usually one or more large segments of their genomes are identical-by-descent (IBD), depending on their degree of relatedness and the recombination and segregation of alleles^12^. Distant cryptic relatedness is usually limited in study cohorts built from small samplings of large populations, but the level of cryptic relatedness increases substantially as the effective population size decreases and the sample size increases. Finally, unless designed to collect families, traditional population-based studies typically have very little *family structure:* the third “class” of relatedness, consisting of first- and second-degree relationships^2,4,13-15^ (Figure 1B, C).

In contrast, the HPG study design enriches for family structure in several ways. First, HPG studies heavily sample from specific healthcare system regions, and the number of pairs of related individuals ascertained increases combinatorially as more individuals are sampled from a single region (Figure 1A). Second, families who live in the same geographic area likely receive medical care from the same doctors at the same healthcare system due to referrals, shared insurance coverage, and convenience. Third, families who have visited a healthcare system for many years with multiple encounters will have extensive medical records, making them more likely to be included in a study compared to transient residents with brief medical records and fewer encounters. Both family structure and distant cryptic relatedness are more pronounced in populations with low migration rates^5^. Conversely, confounding population substructure may be less of a factor in HPG studies if the sampled healthcare system’s population is a single homogenous genetic deme^1^. As a result, we expect to see an enrichment of family structure in HPG studies compared to random ascertainment of a population. In this article, we focus on family structure and its prevalence in a HPG study using both simulated and real data.

The increase in family structure within HPG studies has significant implications when choosing and executing downstream analyses and must be considered thoughtfully^16-22^. Some tools assume all individuals are unrelated (e.g. PCA), some effectively handle estimates of pairwise-wise relationships (e.g. linear mixed models), and others can directly leverage pedigree structures (e.g. linkage and TDT analyses). The following is a description of some common analysis tools and their use cases based on the varying levels of family structure within large population-based datasets (Figure 1D).

Removal of family structure (i.e. selectively excluding samples to eliminate relationships) is a viable option if a dataset has few closely related samples^4,13-15^ and if the size of the unrelated subset is acceptable for the statistical analysis being performed. For example, principal components (PCs) can be computed on an unrelated subset of the data, and then all samples can be projected onto these PCs^1^. A number of methods exist to compute the maximally sized unrelated set of individuals^23,24^. However, this strategy reduces the sample size and power while discarding potentially valuable relationship information. In practice, the degree of information loss is unacceptable for many analyses if the dataset has even a moderate level of family structure.

Several statistical methods have been developed that explicitly model estimates of pairwise relationships. For example, mixed models provide better power for genome-wide association studies and reduce type-one error compared to methods that do not model the confounding relatedness^19,25-27^, but mixed models become computationally expensive when applied at scale. Pairwise relationships can also be used in a pedigree-free QTL linkage analysis^28^. Additional software packages exist that model population structure and family structure for pairwise relationship estimation (PCrelate)^29^ and principal component analysis (PC-AiR)^30^.

Extensive family structure can be used to reconstruct informative pedigree structures directly from the genetic data with tools like PRIMUS^31^ and CLAPPER^32^, opening the door to a number of pedigree-based methods and analyses that are particularly useful for investigating the correlation between rare variation and disease^33^. Methods that use pedigree structures to aid in identifying the genetic cause of a given phenotype typically involve innovative variations on association mapping, linkage analysis, or both, including MORGAN^34^, pVAAST^17^, FBAT (Web Resources), QTDT (Web Resources), ROADTRIPS^35^, rareIBD^36^, and RV-GDT^37^. The appropriate method to use depends on the phenotype, mode of inheritance, ancestral background, pedigree structure/size, number of pedigrees, and size of the unrelated dataset^38^. Genetically reconstructed pedigrees and estimated relationships can be used in a number of other ways beyond association analyses, including pedigree-aware imputation, pedigree-aware phasing, Mendelian error checking, compound heterozygous knockout detection, *de novo* mutation calling, empirical validation of variant calling methods, and rare variant family-based segregation analysis.

We demonstrate the value of identifying family structure in a large clinical cohort as part of the DiscovEHR study. This cohort of 92,455 exomes originated from a collaborative, ongoing study by the Regeneron Genetics Center (RGC) and the Geisinger Health System (GHS) initiated in 2014^1^. DiscovEHR is a dense sample of patient-participants from a single healthcare system that serves a largely rural population in central Pennsylvania with low migration rates. We identify a tremendous amount of family structure within the DiscovEHR cohort, and our simulations project that 70%-80% of the individuals in our sequenced cohort will have a first-or second-degree relative as we continue sequencing up to 250K individuals. This has significant implications on downstream analyses, but also affords us the opportunity to leverage the rich family structure through pedigree reconstruction, phasing compound heterozygous mutation (CHM), and detecting *de novo* mutations (DNM).

## Subjects and Methods

### Patients and samples

We sequenced the exomes of 93,368 de-identified patient-participants from the Geisinger Health System (GHS). Participants were consented into the MyCode® Community Health Initiative^38^ and contributed DNA samples for genomic analysis as part of the Regeneron-GHS DiscovEHR collaboration^1^. Each patient has their exome linked to a corresponding de-identified electronic health record (EHR). A more detailed description of the first 50,726 sequenced individuals has been previously published^1,39^.

The DiscovEHR Study did not specifically target families to participate in the study, but implicitly enriched for adults with chronic health problems who interact frequently with the healthcare system (and may be related to each other), as well as participants from the Coronary Catheterization Laboratory and the Bariatric Service from GHS.

### Sample preparation, sequencing, variant calling, and sample QC

Sample prep and sequencing for the first ∽61K samples have been previously described^1^ and this set of samples are referred to in this manuscript as the “VCRome set”. The remaining set of ∽31K samples were prepared in the same process, except in place of the NimbleGen probed capture we used a slightly modified version of IDT’s xGen probes; supplemental probes were added to capture regions of the genome well-covered by the NimbleGen VCRome capture reagent but poorly covered by the standard xGen probes. Captured fragments were bound to streptavidin-conjugated beads and non-specific DNA fragments were removed by a series of stringent washes according to the manufacturer’s recommended protocol (IDT). We refer to this second set of samples as the “xGen set”. Variant calls were produced using the Genome Analysis Toolkit (GATK; Web Resources). GATK was used to conduct local realignment of the aligned, duplicate-marked reads of each sample around putative indels. GATK’s HaplotypeCaller was then used to process the INDEL-realigned, duplicate-marked reads to identify all exonic positions at which a sample varied from the genome reference in the genomic VCF format (GVCF). Genotyping was accomplished using GATK’s GenotypeGVCFs on each sample and a training set of 50 randomly selected samples outputting a single-sample VCF file identifying both SNVs and indels as compared to the reference. The single sample VCF files were used to create a pseudo-sample that contained all variable sites from the single sample VCF files in both sets. Independent pVCF files were created for the VCRome set by joint calling 200 single-sample gVCF files with the pseudo-sample to force a call or no-call for each sample at all variable sites across the two capture sets. All 200-sample pVCF files were combined to create the VCRome pVCF file. This process was repeated to create the xGen pVCF file. The VCRome and xGen pVCF files were then combined to create the union pVCF. We aligned sequence reads to GRCh38 and annotated variants using Ensembl 85 gene definitions. We restricted the gene definitions to 54,214 transcripts that are protein-coding with an annotated start and stop, corresponding to 19,467 genes. After the previously described sample QC process, 92,455 exomes remained for analysis.

### Principal components and ancestry estimation

We used PLINKv1.9^24^ to merge the union datasets with HapMap3^40^ and kept only SNPs that were in both datasets based on rsID. We also applied the following PLINK filters: --maf 0.1 --geno 0.05 --snps-only --hwe 0.00001 to obtain a set of high-quality common variants. We calculated principal component (PC) analysis for the HapMap3 samples and then projected each sample in our dataset onto those PCs using PLINK. We used the PCs for the HapMap3 samples to train a kernel density estimator (KDE) for each of the five ancestral super classes: African (AFR), admixed American (AMR), east Asian (EAS), European (EUR), and south Asian (SAS). We used the KDEs to calculate the likelihood that each sample belongs to each of the super classes. For each sample, we assigned the ancestral superclass based on the likelihoods. If a sample has two ancestral groups with a likelihood > 0.3, then we assigned AFR over EUR, AMR over EUR, AMR over EAS, SAS over EUR, AMR over AFR; otherwise “UNKNOWN” (this was done to provide stringent estimates of the EUR and EAS populations and inclusive estimates for the more admixed populations in our dataset). If zero or more than two ancestral groups had a high enough likelihood, then the sample was assigned “UNKNOWN” for ancestry. Samples with unknown ancestry were excluded from the ancestry-based identity-by-descent (IBD) calculations.

### IBD estimation

Genome-wide identity-by-descent (IBD) estimates are a metric to quantify the level of relatedness between pairs of individuals^28^. We applied the same Hardy-Weinberg equilibrium, minor allele frequency, and variant level missingness that we applied during the PCA analysis. Next, we used a two-pronged approach to obtain accurate IBD estimates from the DiscovEHR cohort exomes. First, we calculated IBD estimates among individuals within the same ancestral superclass (e.g. AMR, AFR, EAS, EUR, and SAS) as determined from our ancestry analysis. We used the following PLINK flags to obtain IBD estimates out to second-degree relationships: --genome --min 0.1875. This allows for more accurate relationship estimates because all samples share similar ancestral alleles; however, this approach is unable to predict relationships between individuals with different ancestral backgrounds, e.g. a child of a European father and Asian mother.

Second, in order to catch the first-degree relationships between individuals with different ancestries, we calculated IBD estimates among all individuals using the - - min 0.3 PLINK option. We then grouped individuals into first-degree family networks where network nodes are individuals and edges are first-degree relationships. We ran each first-degree family network through the prePRIMUS pipeline^31^, which matches the ancestries of the samples to appropriate ancestral minor allele frequencies to improve IBD estimation. This process accurately estimates first-and second-degree relationships among individuals within each family network (minimum PI_HAT of 0.15).

Finally, we combined the IBD estimates from the two previously described approaches by adding in any missing relationships from family network derived IBD estimates to the ancestry-based IBD estimates. This approach resulted in accurate IBD estimates out to second-degree relationships among all samples of similar ancestry and first-degree relationships among all samples.

IBD proportions for third-degree relatives are challenging to accurately estimate from large exome sequencing dataset with diverse ancestral backgrounds because the analysis often results in an excess number of predicted 3^rd^ degree relationships due to artificially inflated IBD estimates. We used a --min 0.09875 cutoff during the ancestry specific IBD analysis to get a sense of how many third-degree relationships we may have in the DiscovEHR cohort, but these were not used in any of the phasing or pedigree-based analyses. Rather for the relationships-based analyses reported in this paper, we only used high-confidence third-degree relationships we identified within first-and second-degree family networks.

During QC prior to creating the final set of 92,455 individuals, we removed all identical pairs of samples (PI_HAT > 0.9) unless GHS was able to find evidence through a chart review that the two corresponding individuals appear to be different people, share the same birthdate, and had one or more additional pieces of information that they were related (e.g. same last name, shared parent, same address, or listed the other as a relative).

### Pedigree reconstruction

We reconstructed all first-degree family networks identified within the DiscovEHR cohort with PRIMUSv1.9.0^31^. The combined IBD estimates were provided to PRIMUS along with the genetically derived sex and EHR reported age. We specified a relatedness cutoff of PI_HAT > 0.375 to limit the reconstruction to first-degree family networks, and a minimum cutoff of 0.1875 to define second-degree networks.

### Allele-frequency-based phasing

We phased all bi-allelic variants from the VCRome and xGen exome datasets separately using EAGLEv2.3^41^. In order to parallelize our analysis, we divided the genome into overlapping segments of ∽40K variants with a minimum overlap of 500 variants and 250K base-pairs. Since our goal was to phase putative compound heterozygous mutations within genes, we took care to have the segment break points occur in intergenic regions.

We used the UCSC LiftOver program to lift-over EAGLE’s provided genetic_map_hg19.txt.gz file from hg19 to GRCh38 and removed all variants that switched chromosomes or changed relative order within a chromosome resulting in the cM position to not be increasing when sorting on increasing chromosome position. In most cases, this QC step removed inversions around centromeres. We also removed all SNPs that mapped to an alternate chromosome. In total, only 2,783 of the 3.3 million SNPs were removed from the genetic map file. We provided the data for each segment to EAGLE as PLINK formatted files and ran it on DNAnexus with the following EAGLE command line parameters:

- - geneticMapFile=genetic_map_hg19_withX.txt.GRCh38_liftover.txt.gz

- - maxMissingPerIndiv 1

- - genoErrProb 0.01

- - numThreads=16

### Compound heterozygous calling

Our goal was to obtain high confidence compound heterozygous mutation (CHM) calls of putative loss-of-function (pLoF) variants to identify humans with both copies of genes potentially knocked out or disrupted. We classify variants as pLoFs if they result in a frameshift, stop codon gain, stop codon loss, start codon gain, start codon loss, or splicing acceptor or donor altering variant. We created a second, expanded set of potentially harmful variants that included the pLOFs as well as likely disruptive missense variants, which are variants predicted to be deleterious by all five of the following methods: SIFT^41^ (damaging), PolyPhen2 HDIV^42^ (damaging and possibly damaging), PolyPhen2 HVAR (damaging and possibly damaging), LRT^43^ (deleterious), and MutationTaster^44^ (disease causing automatic and disease causing).

We identified rare (alternate allele frequency < 1%) potential compound heterozygous mutations (pCHMs) by testing all possible combinations of heterozygous pLoFs and/or deleterious missense variants within a gene of the same person. We excluded all variants that were out of Hardy-Weinberg equilibrium (p-value < 10^−15^ calculated with PLINKv1.9^24^), that exceeded 10% missingness within the individuals capture specific dataset (i.e. VCRome or xGen sets), or that had another variant within 10 base-pairs in the same individual. We also excluded SNPs with quality by depth (QD) < 3, alternate allele balance (AB) < 15%, and read depth < 7, and we excluded indels with QD < 5, AB < 20%, and read depth < 10. After filtering, we had 57,355 high-quality pCHMs distributed among 36,739 individuals that could knockout or disrupt the function of both copies of a person’s gene if the pCHM variants are phased in *trans*.

The next step was to phase the pCHMs. We used a combination of population allele-frequency-based phasing with EAGLE and pedigree/relationship-based phasing to determine if the pCHMs were in *cis* or *trans*. Figure 2 diagrams the pCHM phasing workflow we employed to obtain the most accurate phasing for each pCHM. Trios and relationships with individuals in both the VCRome and xGen datasets were used only if both variants in the pCHM were on both the VCRome and our modified xGen capture designs. Trio and relationship phasing proved to be more accurate than EAGLE phasing (Table S1), so we preferentially used the pedigree and relationship data for phasing. Table S2 describes our logic used to determine phase of the pCHMs for the different types of familial relationships. For all remaining pCHMs, we used the EAGLE phased data described above. We excluded any EAGLE phased pCHM where one or both of the variants was a singleton because EAGLE phasing accuracy with singletons was not significantly different than random guessing (Table S3). We found that if the two variants in the pCHM have the same minor allele count (MAC) less than 100, then they are in *cis* (22 out of 22 occurrences in child of trios) in our dataset.

**Figure 2.**
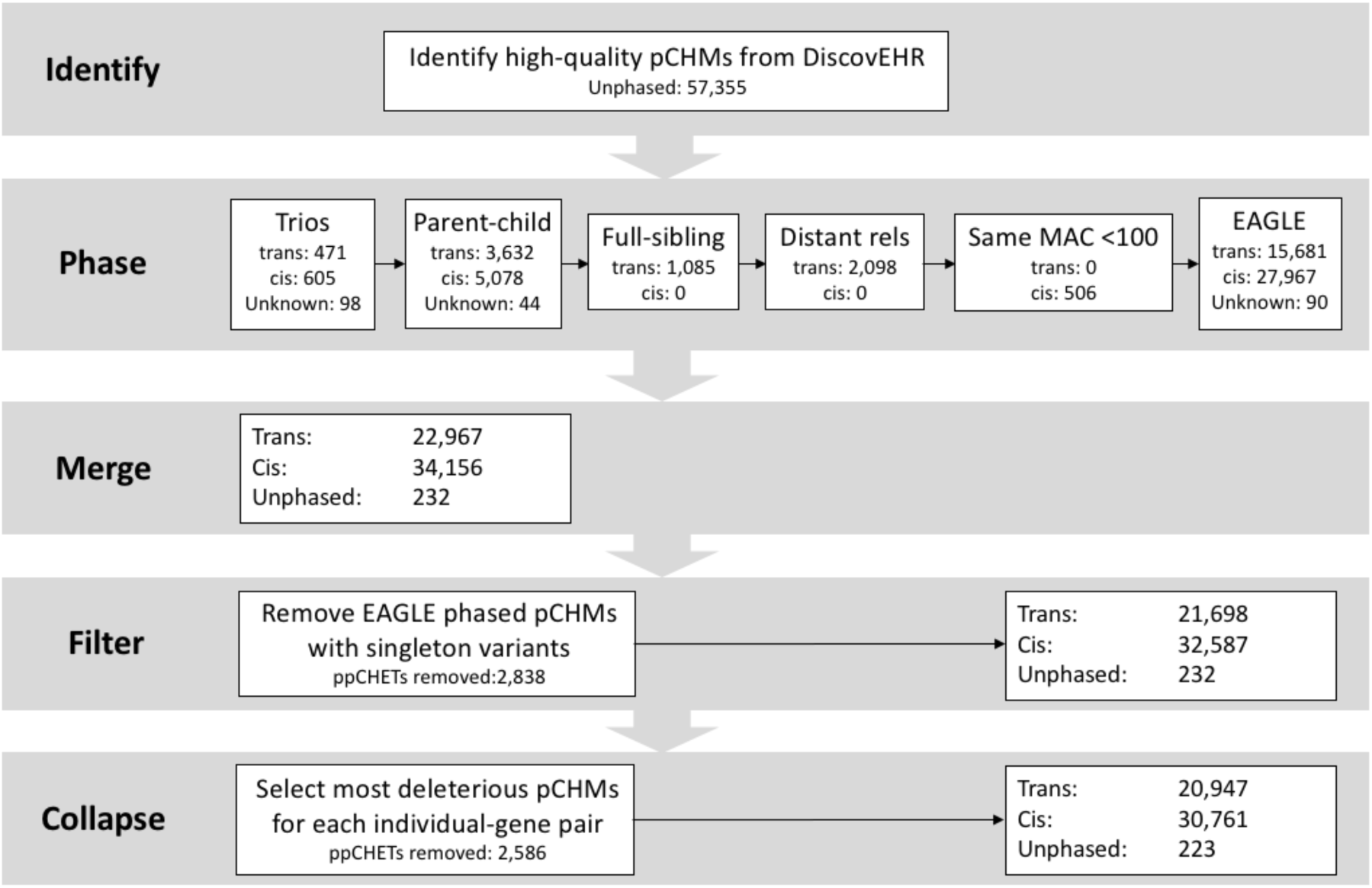
*Decision cascade for determining the phase of potential compound heterozygous mutations (pCHMs) among the 92K DiscovEHR participants. 25.1% of pCHMs and 33.8% of the CHMs (trans) were phased with trios or relationships data.*

We used the trio-phased pCHMs as the truth set to evaluate the overall phasing accuracy of EAGLE. However, if we included the parents of the children of trios in the EAGLE phasing dataset, then EAGLE will use their haplotypes to more accurately phase the children’s variants. Having the parental haplotypes in the dataset improves the phasing accuracy of the children’s pCHMs but does not provide an accurate estimate of phasing accuracy across the entire dataset. To obtain a good measure of accuracy for the EAGLE pCHM phasing across the entire cohort, we reran EAGLE on the entire dataset as before but excluded all first-degree relatives of one child in each nuclear family before phasing. We then compared the EAGLE phased pCHMs to the trio-phased pCHMs to estimate the overall EAGLE phasing accuracy.

Finally, if there were more than one pCHM within the same gene of an individual, then only the pCHM with the most deleterious profile was retained (Table S4). Using the approach outlined above, we were able to phase >99% of all pCHMs, and identify 20,947 rare compound heterozygous mutations (CHMs) that are predicted to be function altering.

## Compound heterozygous mutation validation

We evaluated phasing accuracy by comparing phasing predictions to phasing done with trios and with Illumina reads. We performed Sanger validation on a subset of the incorrectly phased pCHMs to see if the variants were false positive calls.

First, phasing accuracy of the pCHMs was evaluated by using the trio phased pCHMs as truth. Since the phasing approach of each familial relationship is performed independently from the trio phasing, we can get a good measure of phasing accuracy of each of the relationship classes as long as the pCHM carrier is a child in a trio. Table S1 shows that the accuracy of family-based phasing was 99.6% (1060/1064 pCHMs) for rare pCHMs. EAGLE phasing was less accurate at 89.1% (766/860 pCHMs; Table S1). We evaluated the accuracy of EAGLE at phasing pCHMs in different minor allele frequency ranges, and found that it consistently attains an accuracy greater than 90% with a MAC greater than 9 and ∽77% for a MAC between 2-9 (Table S3). EAGLE phasing performed poorly with singletons.

Second, we attempted to validate 200 pCHMs with short Illumina reads (∽75 bp) by looking at the read stacks in the Integrative Genomics Viewer (IGV)^45^ to see if the two variants occur on the same read or independently. We were able to decisively phase 190 (115 *cis* and 79 *trans*; 126 EAGLE phased and 74 pedigree/relationship phased) selected pCHMs using short reads. The remaining ten showed read evidence of both *cis* and *trans* phasing, most likely due to one or both of the variants being a false positive call. Visual validation showed an overall accuracy of 95.8% and 89.9% for pedigree/relationships and EAGLE phasing, respectively (Table S5). While the Illumina read-based validation results are in line with the trio validation results, we do note that the Illumina read-based validation accuracy results are lower than the phasing accuracy determined by phasing with trios. The difference is likely due to the enrichment for false positive pCHMs in small problematic regions of exons prone to sequencing and variant calling errors.

### *De novo* mutation (DNM) detection

We merged the results from two different approaches for detecting DNMs. The first method is TrioDeNovo^46^, which reads in the child’s and parents’ genotype likelihoods at each of the child’s variable sites. These likelihoods are input into a Bayesian framework to calculate a posterior likelihood that a child’s variant is a DNM. The second program is DeNovoCheck (Web Resources), which is described in the supplemental methods of de Ligt, et al.^47^. DeNovoCheck takes in a set of candidate DNMs identified as being called in the child and not in either parent. It then verifies the presence of the variant in the child and absence in both of the parents by examining the BAM files. We filter these potential DNMs and evaluate a confidence level for each DNM in the union set using a variety of QC metrics. Figure S1 illustrates this DNM calling process, shows the variant filters we applied, and provides the criteria we used to classify each DNM as either low-confidence, moderate-confidence, or high-confidence. We excluded all low-confidence and non-exonic DNMs from the summary results of this paper, but we considered them when doing visual validation to estimate the false negative rate of excluding them. We also excluded the DNM calls for one extreme outlying participant who had an order of magnitude more DNMs called than any other sample.

### LDLR tandem duplication distant pedigree estimation

Although we cannot know the true family history of the de-identified individuals in our cohort, we have used PRIMUS^31^ reconstructed pedigrees, ERSA^12^ distant relationship estimate, and PADRE^48^ to connect the pedigrees to identify the best pedigree representation of the mutation carriers of a novel tandem duplication in LDLR^49^. We used HumanOmniExpress array data (available for 25 out of the 37 carriers) to estimate the more distant relationships and used the method as described in the PADRE to connected the PRIMUS reconstructed pedigrees.

### SimProgeny

We developed a forward simulation framework (SimProgeny) to simulate a wide variety of populations, including a population served by a healthcare system like GHS. SimProgeny also simulates sample ascertainment used by HPG studies (Figure S2). SimProgeny can simulate populations of millions of people dispersed across one or more sub-populations based on user specified population parameters (Table S6). The simulation progresses year-to-year simulating couplings, births, separations, migrations, deaths, and movement between sub-populations based on specified parameters. This process generates realistic pedigree structures and populations that represent a wide variety of HPG studies. The default values have been tuned to model the DiscovEHR cohort, but these parameters can be easily customized to model different populations by modifying the configuration file included with the SimProgeny code available at (Web Resources). See Supplemental Methods for a detailed description of SimProgeny.

In addition to modeling populations, SimProgeny simulates two ascertainment approaches to model selecting individuals from a population for a genetic study: random ascertainment and clustered sampling. Random ascertainment gives each individual in the population an equal chance of being ascertained without replacement. Clustered sampling is an approach to enrich for close relatives, and it is done by selecting an individual at random along with a number of their first-and second-degree relatives. The number of first-degree relatives is determined by sampling a value from a Poisson distribution with a user specified first-degree ascertainment lambda (default is 0.2). The number of second-degree relatives is determined in the same way and the default second-degree ascertainment lambda is 0.03. See Supplemental Methods for additional information on SimProgeny’s ascertainment options.

### Simulation of the underlying DiscovEHR population and its ascertainment

Our DiscovEHR simulations contained individual populations with starting sizes of 200K, 300K, 350K, 400K,475K, 500K, and 550K. We tuned the SimProgeny parameters (Table S6) with publically available country, state, and county level data as well as our own understanding of how individuals were ascertained through GHS consenting and sample collection. Sources for the selected parameters are available in supplemental file Simulation_parameters.xls. We reduced the immigration and emigration rates from the state-wide Pennsylvania (PA) average given that GHS primarily serves rural areas that tend to have lower migration rates than more urban areas. Simulations were run with a burn-in period of 120 years and then progressed for 101 years. Simulated populations grew by ∽15%, which is similar to the growth of PA since the mid-20^th^ century.

We performed both random and clustered ascertainment. For both ascertainment approaches, we shuffled the ascertainment order of the first 5% of the population (specified with the ordered_sampling_proportion parameter) to model the random sequencing order of the individuals in GHS biobank at the beginning of our collaboration. While the selection of this parameter has no effect on random ascertainment and a negligible effect on the accumulation of pairwise relationships in clustered ascertainment, it does affect the proportion of individuals with one or more relatives in the clustered sampling dataset by creating an inflection point at 5% population ascertainment in the simulation results plots (Figure S3B, D). This inflection point would be less pronounced if we were to model the freeze process of the real data or model a smoother transition between sequencing samples from the biobank and newly ascertained individuals. Notably, the inflection point is more pronounced with higher values of lambda from the Poisson distribution.

## Results

### Relationship estimation and relatedness in DiscovEHR

In the current dataset of 92,455 individuals, we identified 43 monozygotic twins, 16,476 parent-child relationships, 10,479 full-sibling relationships, and ∽39,000 second-degree relationships (Figure 3A). Next, we treated individuals as nodes and relationships as edges to generate undirected graphs. Using only first-degree relationships, we identified 7,684 connected components, which we refer to as first-degree family networks. Figure 3B shows the distribution in size of the first-degree family networks, which range from 2 to 25 sequenced individuals. Similarly, we found 10,173 second-degree family networks; the largest containing 19,968 individuals (∽22% of the overall dataset; Figure 3C). We were able to identify ∽5,300 third-degree relationships within the second-degree family networks. Using a lower IBD cutoff (PI_HAT > 0.09875) for the IBD estimations within ancestral groups without consideration of second-degree family networks, we found well over 100,000 third-degree relationships within the DiscovEHR cohort. Given that 95.9% of DiscovEHR individuals are of European ancestry (Table S7), it is not surprising that the vast majority (98.6%) of the pairwise relationships found are between two individuals of European ancestry (Table S8). Nonetheless, we identified many relationships between people of the same, non-European ancestry and between individuals with different ancestries; for example, there are several trios having one European parent, one East Asian parent, and a child whose ancestry is unassigned to a super-population given the ad-mixed nature of his/her genome.

**Figure 3.**
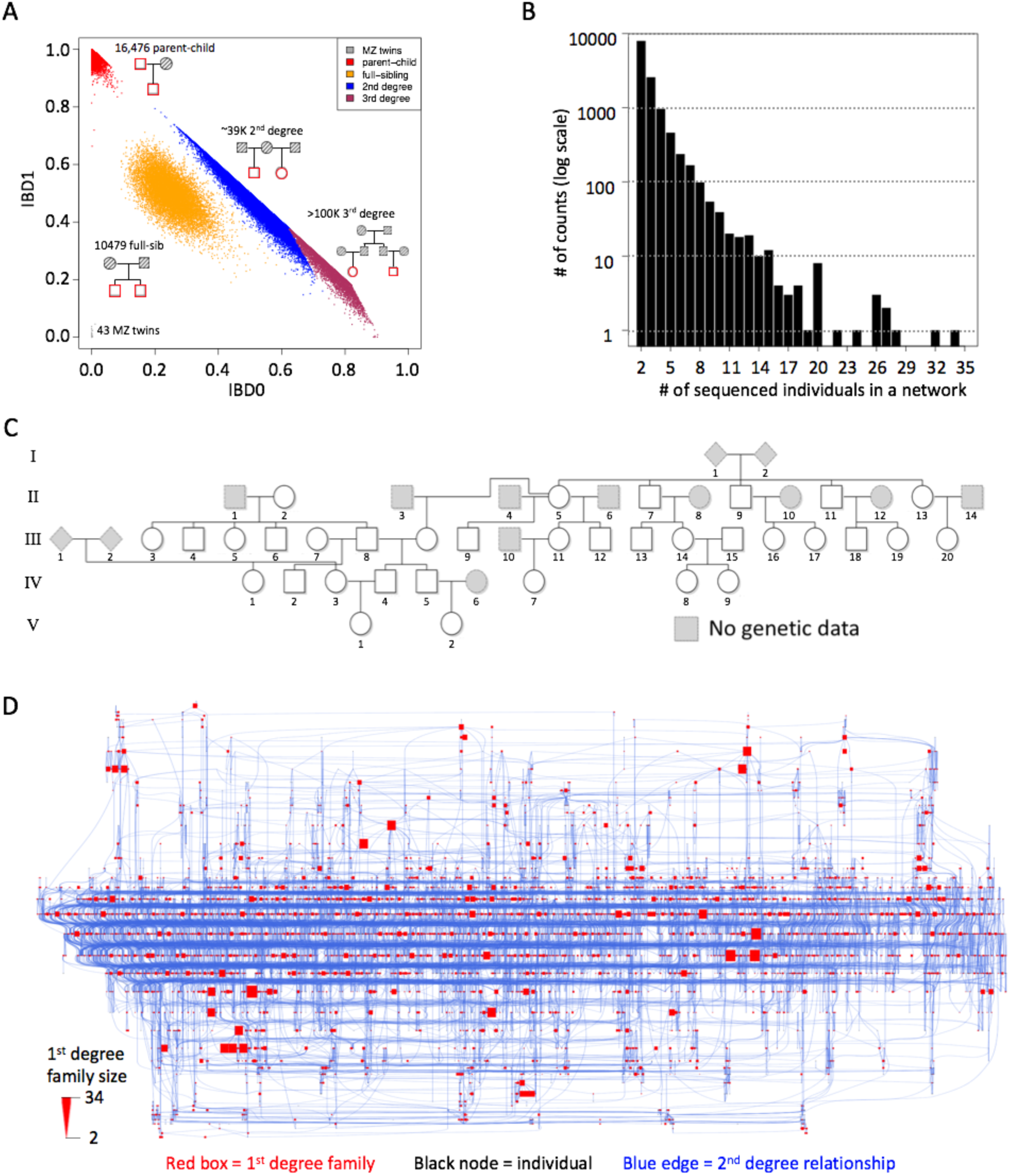
*First 92K sequenced individuals from the DiscovEHR cohort contain an extensive amount of relatedness. (A) IBD0 vs IBD1 plot shows pairwise relationships segregating into different familial relationship classes. The IBD sharing distributions of second-and third-degree relationships overlap with each other, so a hard cutoff halfway between the two expected means was selected. (B) The distribution of size of first-degree family networks ranges between 2 and 34 sequenced individuals, with a vast majority of smaller family networks. (C) More than 99.98% of the first-degree family networks’ pedigree structures were reconstructed using the pairwise-IBD estimates, including this pedigree of 34 sequenced individuals. (D) The largest second-degree family network of 19,968 (∽22% of the dataset) shows 4,062 first-degree family networks (red boxes proportionally sized to the number of individuals in the network; including the pedigree shown in C) and 5,584 additional individuals (black nodes) connected by 11,430 second-degree relationships (blue edges). ^*\**^Third-degree relationships are challenging to accurately estimate due to technical limitations of exome data as well as the widening and overlapping variation around the expected mean IBD proportions of more distant relationship classes (e.g. fourth-degree and fifth degree). We provided a lower bound estimate of the number of third-degree relationships.*

Importantly, we show both empirically (Figure 4A) and through simulation (Figure 5) that the rate of accumulating relatives far exceeds the rate of ascertaining samples. This is expected since there are combinatorially increasing numbers of possible pairwise relationships within the dataset as the size increases, and the likelihood that a previously unrelated individual in the dataset becomes involved in a newly identified relationship also increases. Currently, 39% of individuals in the DiscovEHR cohort have at least one first-degree relative in the dataset, and 56% of the participants have one or more first-or second-degree relatives in the dataset (Figure 4B).

**Figure 4.**
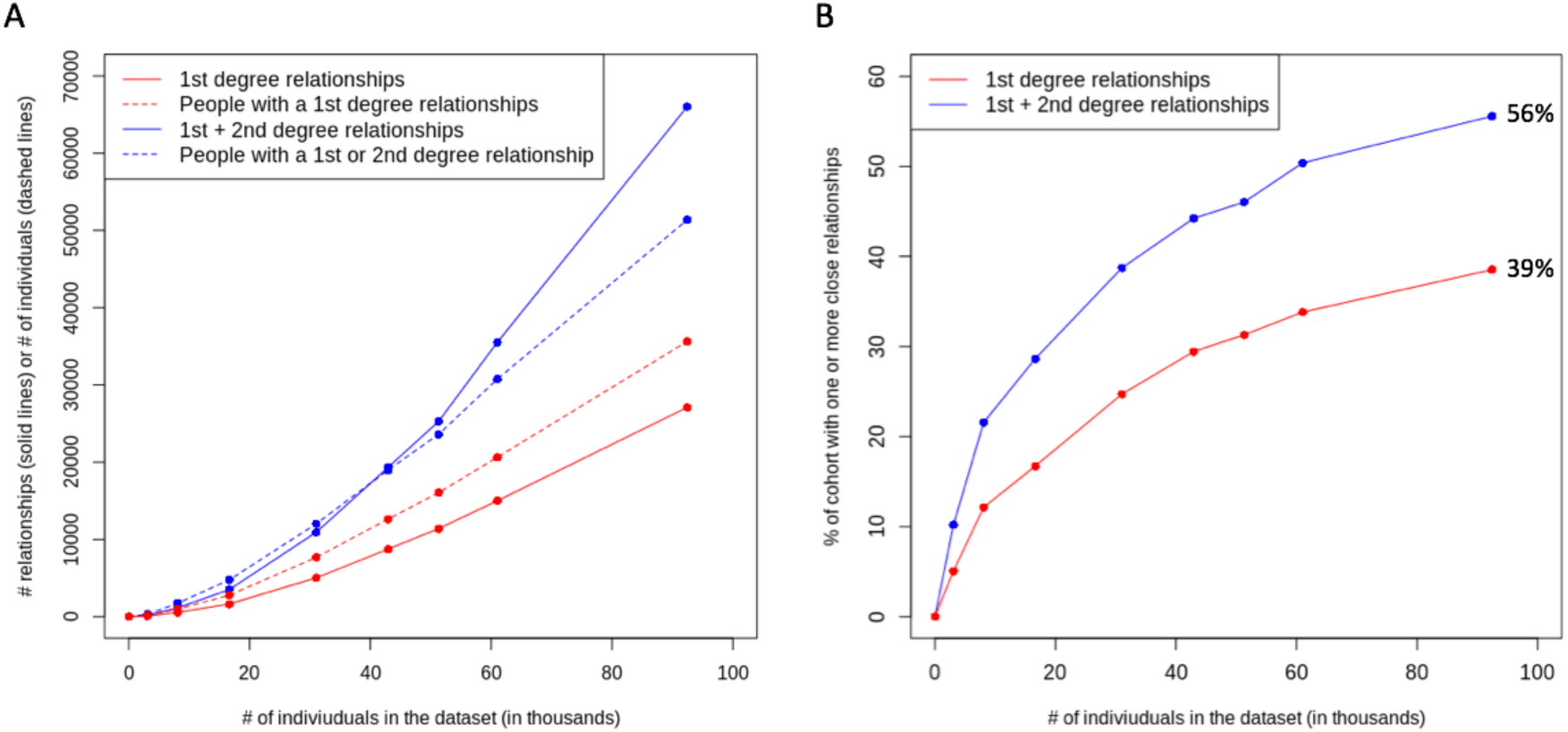
*Accumulation of relatedness within the DiscovEHR cohort at consecutive data freezes. A) The number of pairwise relationships has grown rapidly. B) The proportion of individuals in the cohort having a first-or second-degree relative identified in the cohort.*

**Figure 5.**
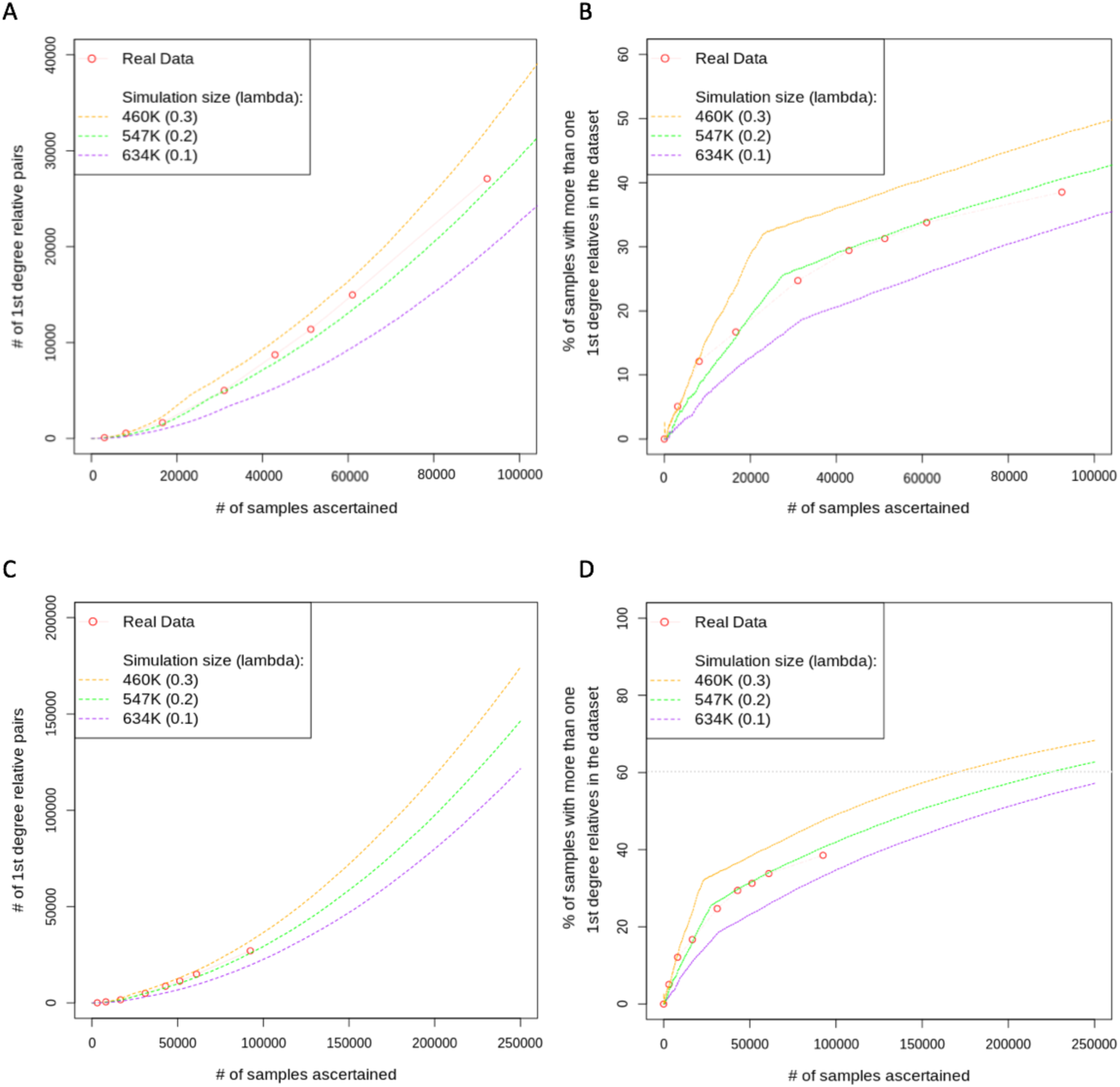
*Simulated population and ascertainment fit to the accumulation of first-degree relatedness in the DiscovEHR cohort. The real data was calculated at periodic “freezes” indicated with the punctuation points connected by the faint red line. Most simulation parameters were set based on information about the real population demographics and the DiscovEHR ascertainment approach. However, two parameters were unknown and selected based on fit to the real data: 1. the effective population size from which samples were ascertained and 2. the increased chance that someone is ascertained given a first-degree relative previously ascertained, which we call “clustered ascertainment”. All panels show the same three simulated population sizes spanning the estimated effective population size. We simulated clustered ascertainment by randomly ascertaining an individual along with a Poisson-distributed random number of 1^st^ degree relatives (distributions’ lambdas are indicated in the legends). (A) The accumulation of pairs of first-degree relatives as additional samples are ascertained. (B) The proportion of the ascertained participants that have one or more first-degree relatives that have also been ascertained. (C) Simulated ascertainment projections with upper and lower bounds of the number of first-degree relationships we expect with our current DiscovEHR ascertainment approach as we scale to our goal of 250K participants. (D) Simulated projections with upper and lower bounds of the proportion of the ascertained participants that have 1 or more first-degree relatives that have also been ascertained.*

### Simulations with SimProgeny and relatedness projections

Prior to the launch of the DiscovEHR collaboration, it was unclear how much relatedness we should expect to see and whether it would follow the levels of relatedness seen in previous population-based genomic studies. However, it became clear early on that the cohort contained far more family structure than typically seen in population-based studies, and projections estimated that the proportion of the cohort involved in close relationships would eventually involve the majority of our dataset. Given the impact of this relatedness on downstream analyses, we set out to determine whether this amount of relatedness is expected, whether it is unique to our dataset, and how much it would grow as the sequenced cohort expands.

To answer these questions, we developed a flexible simulation framework (SimProgeny) to model a wide variety of study populations and sampling approaches to estimate the amount of relatedness researchers should expect to find for a given set of populations and sampling parameters. While we apply this framework to the DiscovEHR cohort, it is flexible enough that it also can be applied to modeling shallower ascertainment of more transient populations.

We used SimProgeny to simulate the DiscovEHR population and the ascertainment of the first 92,455 participants. As expected, the simulations show that DiscovEHR participants were not randomly sampled from the population, but rather the dataset is enriched for close relatives (Figure S4). Therefore, we used a clustered ascertainment approach (see Methods) that more accurately models ascertainment from a healthcare system study population and the subsequent enrichment of close relatives observed in the real data (Figure 5). These simulation results suggest that the effective population size for the first 60K participants was ∽475K individuals, and a Poisson distribution having lambda of 0.2 most closely matches the enrichment of first-degree relatives. However, the departure of the real data line (Figure 4, faint red line) from the ∽475K simulation line (solid green line) at 90K ascertained samples suggests that the DiscovEHR cohort’s effective population size may have increased after ascertaining the first 60K samples. These estimates are consistent with our knowledge that the majority of the first 30K-60K DiscovEHR participants reside in the counties surrounding the GHS headquarters in Danville, and the participant base subsequently expanded to more heavily include pockets of individuals from north-central and northeast rural Pennsylvania (Figure S5). Most notably, ascertainment was not evenly distributed across the entire GHS catchment area (>2.5 million individuals).

After identifying simulation parameters that reasonably fit the real data, we used SimProgeny to obtain a projection of the amount of first degree relationships we should expect as DiscovEHR expands to our goal of 250K participants. If we continue to ascertain participants in the same way, we expect to obtain ∽150K first-degree relationships (Figure 5C) involving ∽60% of DiscovEHR participants (Figure 5D). We then expanded our simulation analysis to include second-degree relationships, and the simulation results suggest that with 250K participants we should expect well over 200K combined first-and second-degree relationships involving over 70% of the individuals in DiscovEHR (Figure S3).

These projections of relatedness in DiscovEHR assume that we continue ascertaining participants in the same way we did for the first 60K-90K participants. However, if DiscovEHR expands ascertainment of participants to additional GHS clinics and hospitals in other regions, then these relatedness estimates are likely to drop, because expanding the participant base increases the size of the effective sampling population and taps into new genetic demes or distant branches of the same demes. The level of the relatedness will depend on the proportion of the total population we ascertain and the underlying population demographics of the regions, both of which can be simulated with SimProgeny.

While SimProgeny is designed to reasonably model a real population, it has its limitations. For example, the Poisson distribution accurately models the clustered sampling of first-degree relationships we observe in our real data, but it underestimates the clustering of second-degree relationships compared to what is observed in DiscovEHR. Thus, there are bound to be nuances that cannot be easily modeled with a fixed distribution, and there are likely to be other confounding aspects to how participants were ascertained in the real dataset.

Regardless, our simulation results demonstrate a clear enrichment of relatedness in the DiscovEHR HPG study as well as provide key insights into the tremendous amount of relatedness we expect to see as we continue to ascertain additional participants, assuming future ascertainment is reasonably well modeled by SimProgeny. These observations can also be extrapolated to other large HPG studies, and the flexibility built into the model provides the ability to tune the model to a wide variety of different populations and ascertainment approaches.

### Leveraging relatedness instead of treating it like a nuisance

We have reconstructed pedigree structures for 12,574 first-degree family networks in the DiscovEHR data set using the pedigree reconstruction tool PRIMUS^31^, and found that 98.9% of these pedigrees reconstructed unambiguously to a single pedigree structure when considering IBD estimates and reported participant ages. These pedigrees include 2,192 nuclear families (1,841 trios, 297 quartets, 50 quintets, 3 sextets, and 1 septet). Table S9 shows a breakdown of the trios by ancestry. Figure 3C shows the largest first-degree pedigrees, which contains 34 sequenced individuals. We have used these relationships and pedigrees in several ways, and we highlight three main applications in this section.

### Compound Heterozygous mutations

A major goal of human genetics is to better understand the function of every gene in the human genome. Homozygous loss-of-function mutations (LoFs) are a powerful tool to gain insight into gene function by analyzing the phenotypic effects of these “human knockouts” (KOs). Rare (MAF < 1%) homozygous LoFs have been highlighted in recent large-scale sequencing studies and have been critical in identifying many gene-phenotype interactions^1,4,50,51^. While rare compound heterozygous mutations (CHMs) of two heterozygous LoFs are functionally equivalent to rare homozygous KOs, they are more difficult to identify (particularly with short-read sequencing) and are rarely interrogated in large sequencing studies^1,4,50^.

We performed a survey of rare CHMs in the DiscovEHR cohort. First, we identified 57,355 high-quality potential CHMs (pCHMs) consisting of pairs of rare heterozygous variants that are either putative LoFs (pLoF, i.e., nonsense, frameshift, or splice-site mutations) or missense variants with strong evidence of being deleterious (see Methods). Second, we phased the pCHMs using a combination of allele-frequency-based phasing using EAGLE and pedigree-based phasing using the reconstructed pedigrees and relationship data (Figure 2). EAGLE phased the pCHMs with an average of 89.1% accuracy based on trio validation (Table S1). However, because we had extensive pedigree and relationship data within this cohort, we were able to use it to phase 25.2% of the pCHMs and 33.8% of the *trans* CHMs with highly accurate trio and relationship phasing data (≥98.0%; Table S1), reducing inaccurate phasing of *trans* CHMs by approximately a third. The phased pCHMs spanned the entire frequency range from singletons to 1% MAF (Table S10).

After processing, 40.3% of the pCHMs were phased in *trans*, yielding a high-confidence set of 20,947 rare, deleterious CHMs distributed among 17,533 of the 92K individuals (mean = 0.23 per person; max = 10 per person; Figure 6A). The median genomic distance between pCHM variants in *cis* (5,955 bps) was a little more than half the median distance between the pCHMs variants in *trans* (11,600 bps; Figure S6). Nearly a third of the CHMs involved at least one pLoF and 8.9% of the of CHMs consisted of two pLoF variants (Table S11). Over 4,216 of the 19,467 targeted genes contain one or more CHM carriers (Table S12), and 2,468 have more than one carrier (Figure 6B). The ten genes with more than 125 CHM carriers are estimated to be among the most LoF tolerant in the genome based on ExAC pLI scores^4^ (Table S13), so it is no surprise that these genes would contain a higher number of CHMs.

**Figure 6.**
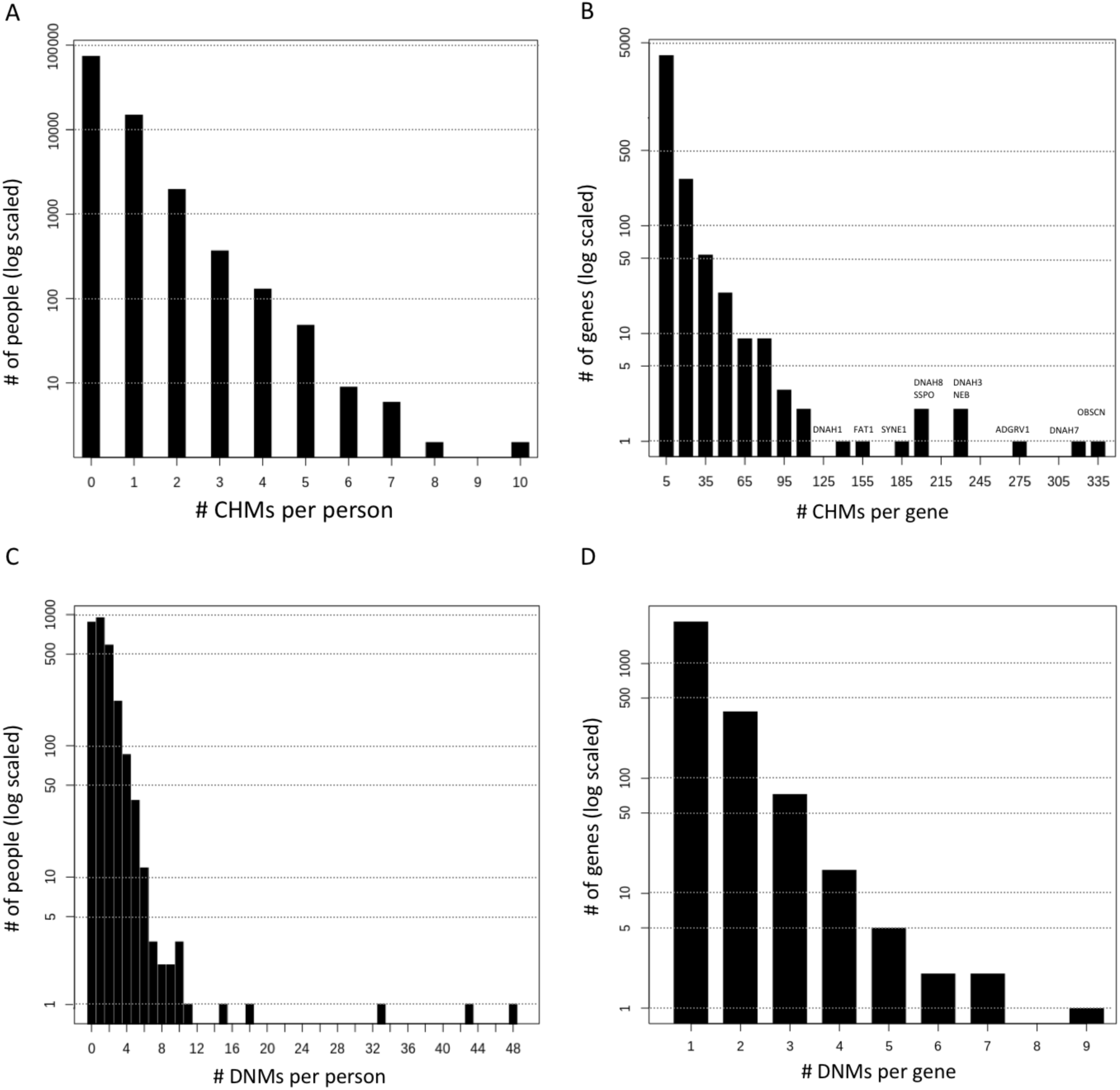
*DiscovEHR results for compound heterozygous mutations (CHMs) and de novo mutations (DNMs). (A) Distribution of the number of CHMs per individual in the DiscovEHR cohort. (B) Distribution of the # of CHMs per gene. Names of genes with more than 125 CHMs are listed. (C) Distribution of 3,415 exonic high and moderate confidence DNMs among the children of trios in the DiscovEHR cohort. (D) The distribution non-synonymous DNMs across the 2,802 genes with 1 or more.*

In order to get a more robust set of human knockout genes and demonstrate the added value of CHMs, we combined the CHMs with the 6,560 rare (MAF < 1%) homozygous pLoFs found among the 92K DiscovEHR participants. pLoF-pLoF CHMs increased the number of genes with ≥1 and ≥20 individuals with a putative KO by 15% and 61%, respectively (Table S12). The benefit of including CHMs in a KO analysis is even more significant when we consider missense variants that are predicted to disrupt protein function. We found a combined 20,364 rare homozygous pLOF and deleterious missense variants among the 92K participants. CHMs provided 26% more genes with ≥1 carriers and 397% more genes with ≥20 carriers where both copies of the gene are predicted to be completely knocked out or disrupted (Table S12).

### De novo mutations

*De novo* mutations (DNMs) are a class of rare variation that is more likely to produce extreme phenotypes in humans due to sporadic occurrence and lack of purifying selection. Many recent sequencing studies have shown that DNMs are a major driver in human genetic disease^47,53,54^, demonstrating that DNMs are a valuable tool to better understand gene function.

We used the nuclear families reconstructed from the 92K DiscovEHR participants to confidently call 3,415 moderate-and high-confidence exonic DNMs distributed among 1,783 of the 2,602 available children in trios (mean = 1.31; max = 48; Figure 6C). PolyPhen2 predicts 29.1% (N=995) of the DNMs as “probably damaging” and an additional 9.2% (N=316) as possibly damaging. The DNMs are distributed across 2,802 genes (Figure 6D) with *TTN* receiving the most at nine. The most common type of DNM is nonsynonymous SNVs (58.5%) followed by synonymous SNVs (24.3%). Table 1 provides a complete breakdown of DNM types and shows that our proportions of DNMs falling into the different functional classes generally match those found in a recent study of DNMs in children with development disorders^53^. We also observed an increase in the number of exonic DNMs with respect to both maternal (0.011 DNMs/year, p=7.3x10^-^4; Poisson regression; Figure S7) and paternal age at birth (0.010 DNMs/year; p=5.6x10^-^4), consistent with other reports^53,55-57^. Notably, maternal and paternal age at birth are highly correlated in our dataset (rho=0.79; Figure S8), thus the rates are not additive and no significant difference was identified to distinguish either as a driving factor (see Supplemental Methods).

**Table 1.**
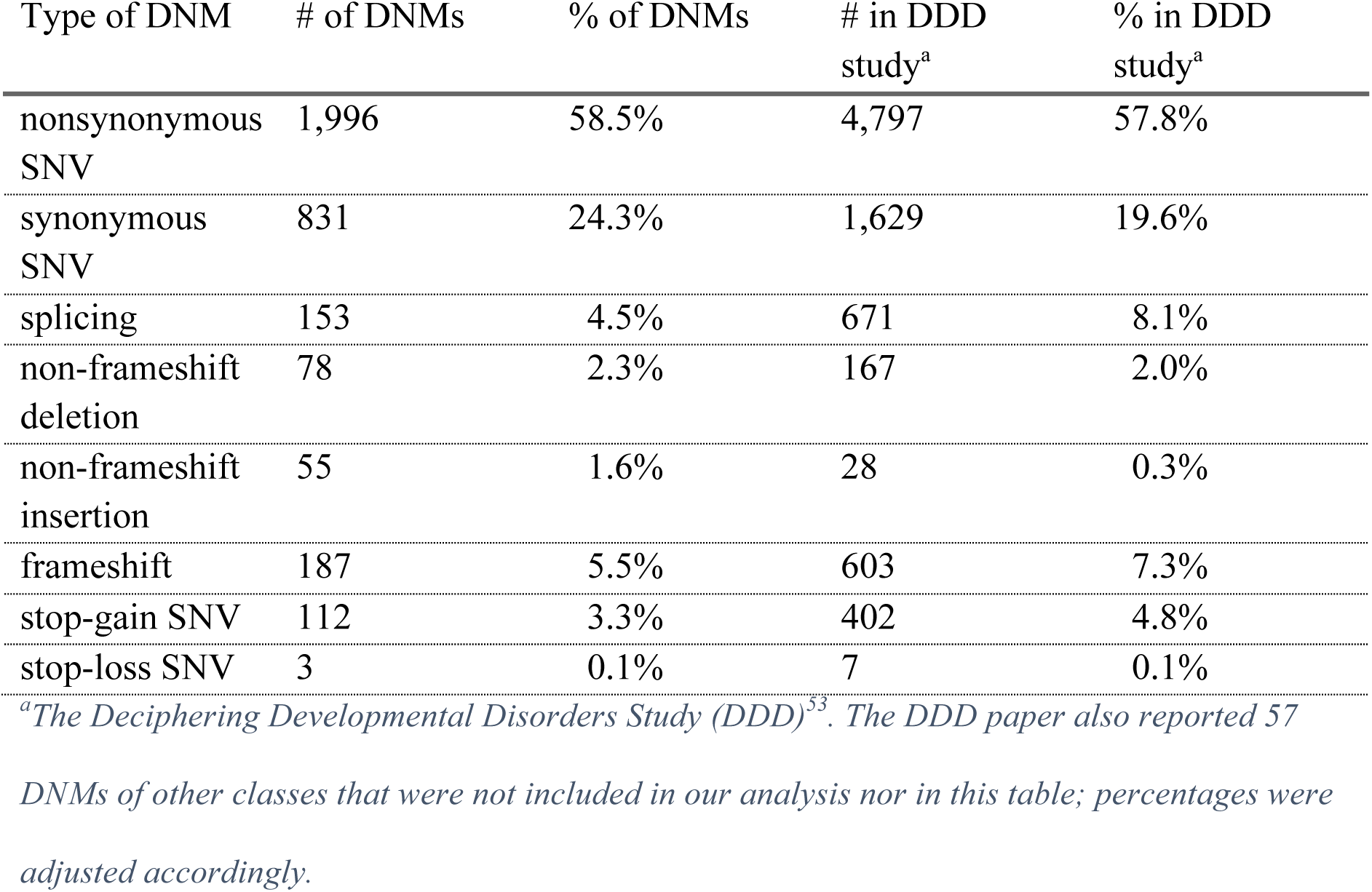
Breakdown by functional class of moderate-and high-confidence exonic DNMs found in the DiscovEHR cohort compared to a recent developmental delay exome study of 4,293 trios.

| Type of DNM | # of DNMs | % of DNMs | # in DDD study<sup>a</sup> | % in DDD study<sup>a</sup> |
| --- | --- | --- | --- | --- |
| nonsynonymous SNV | 1,996 | 58.5% | 4,797 | 57.8% |
| synonymous SNV | 831 | 24.3% | 1,629 | 19.6% |
| splicing | 153 | 4.5% | 671 | 8.1% |
| non-frameshift deletion | 78 | 2.3% | 167 | 2.0% |
| non-frameshift insertion | 55 | 1.6% | 28 | 0.3% |
| frameshift | 187 | 5.5% | 603 | 7.3% |
| stop-gain SNV | 112 | 3.3% | 402 | 4.8% |
| stop-loss SNV | 3 | 0.1% | 7 | 0.1% |

<sup>a</sup>The Deciphering Developmental Disorders Study (DDD)<sup>33</sup>. The DDD paper also reported 57 DNMs of other classes that were not included in our analysis nor in this table; percentages were adjusted accordingly.

We attempted to perform visual validation of 23 high-and 30 moderate-and 47 low-confidence DNMs spanning all functional classes. Eight moderate - and two low-confidence variants could not be confidently called as true or false positive DNMs. Of those remaining, 23/23 (100%) high-confidence, 19/22 (86%) moderate-confidence, and 12/43 (28%) low-confidence DNMs validated as true positives. Visual validation also confirmed that the majority (40/49) of potential DNM in individuals with >10 DNMs are likely false positive calls.

### Variant and phenotype segregation in pedigrees

We have used the reconstructed pedigree data from among the 92K DiscovEHR participants to distinguish between novel/rare population variation and familial variants and have leveraged it to identify highly penetrant disease variants segregating in families. While this is not intended to be a survey of all known Mendelian disease-causing variation transmitted through these pedigrees, we have identified a few illustrative examples including familial aortic aneurysms (Figure 7A), long QT syndrome (Figure 7B), thyroid cancer (Figure 7C), and familial hypercholesterolemia (FH; Figure 8)^49^. The FH example is particularly interesting as we previously reported a novel FH-causing tandem duplication in *LDLR*^49^. We have updated the CNV calls and found 37 carriers of the FH-causing tandem duplication among the 92K exomes, and we have reconstructed 30 out of the 37 carriers into a single extended pedigree. The carriers’ shared ancestral history provides evidence that they all inherited this duplication event from a common ancestor approximately six generations back. While two of the seven remaining carriers are second-degree relatives to each other, genotyping array data was not available to confirm that the remaining seven carriers are also distantly related to the other carriers in Figure 8.

**Figure 7.**
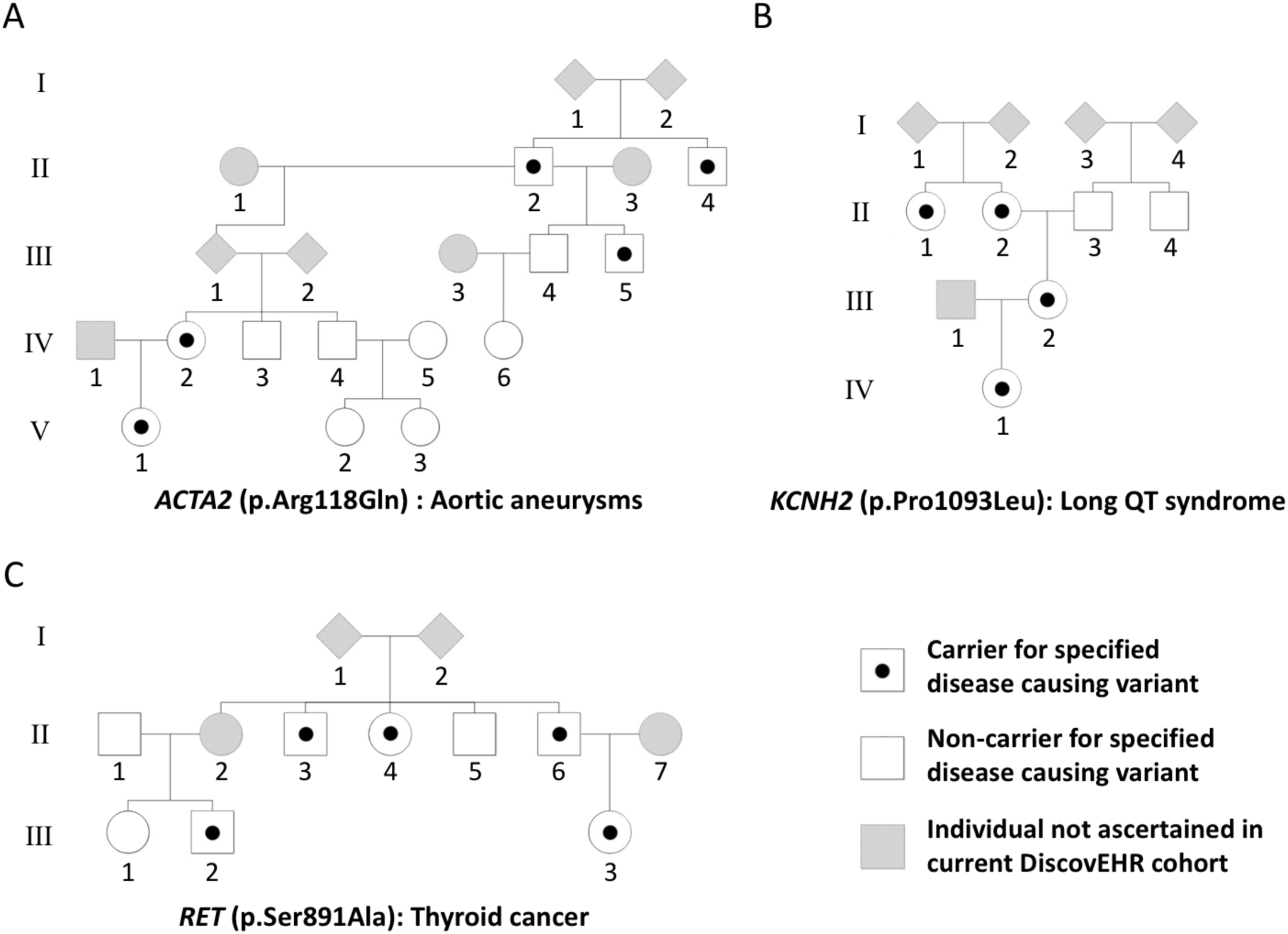
*Reconstructed pedigree from DiscovEHR demonstrating the segregation of known disease-causing variants, including variants for (A) aortic aneurysms, (B) long QT syndrome, and (C) thyroid cancer.*

**Figure 8.**
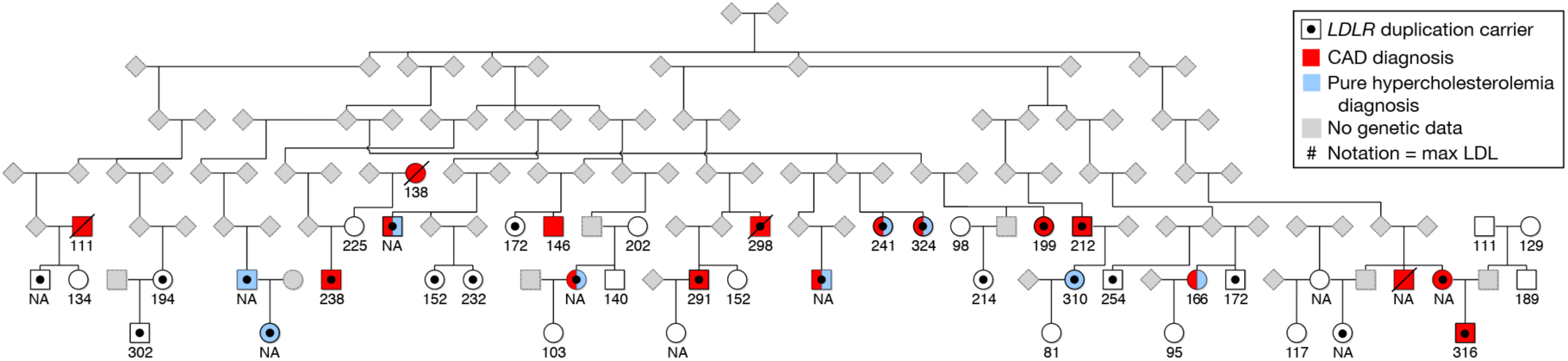
*Image of the reconstructed pedigree prediction containing 25/37 carriers of the novel FH-causing tandem duplication in LDLR and 20 non-carrier, related (first-or second-degree) individuals from the sequenced cohort. Carrier and non-carrier status was determined from the exome data from each individual. Elevated max LDL levels (value under symbols) as well as increased prevalence of coronary artery disease (CAD, red fill) and pure hypercholesterolemia (ICD 272.0; blue) segregate with duplication carriers. Five additional carriers (not drawn) were found to be distant relatives (seventh-to ninth-degree relatives) of individuals in this pedigree.*

## Discussion

Sequencing studies continue to collect and sequence increasing proportions of human populations and are uncovering the extremely complex, intertwined nature of human relatedness. In the first 92K sequenced participants of the DiscovEHR cohort, we have identified ∽66K first-and second-degree relationships, reconstructed 12,574 pedigrees, and uncovered a second-degree family network of nearly 20,000 participants. Studies in founder populations have already highlighted the complexity of relationships (Old Order Amish^58^, Hutterites^59^, and Ashkenazi Jews^60^), and recent studies of non-founder populations are reporting extensive levels of family structure (UK Biobank^61^, NHANES^62^, and AncestryDNA^6^). We observed family structure (first-and second-degree relationships) involving 55.6% of the DiscovEHR participants, and we expect family structure will involve a large proportion, if not a majority, of individuals in other large HPG studies. We have demonstrated through simulations and observations within our own data that we can obtain a large number of close familial relationships, nuclear families, and informative pedigrees within HPG studies. While underlying population structure and depth of ascertainment will vary between studies, we do believe that our observations in DiscovEHR will be applicable to other HPG studies since families tend to visit the same healthcare system and have similar genetic and environmental disease risks. The days of only having a handful of closely related individuals or samples in large sequencing cohorts are over, and we can no longer simply remove closely related pairs of individuals for our association studies knowing that it is only a small fraction of the overall cohort. Instead, we need to continue developing methods that are capable of leveraging the extensive relatedness of these rich cohorts and that can scale to accommodate growing HPG study population sizes and phenotype diversity.

In this study, we have demonstrated several ways to leverage family structure. First, we improved the phasing accuracy of rare compound heterozygous mutations (CHMs). While we did obtain accurate phasing of CHMs with EAGLE, our pedigree- and relationship-based phasing was far more accurate, reducing the pCHM phasing error by approximately a third. We expect that the accuracy of the relationship-based phasing of pCHMs will be lower for variants with >1% MAF because phasing using the pairwise relationships assumes that if two variants appear together in two relatives, then they are in *cis* and have segregated together from a common ancestor. There is a higher chance that two independently segregating common variants will appear together in multiple people, resulting in being incorrectly phased as *cis* by the algorithm. Therefore, common variants may be better phased using population allele frequencies with programs like EAGLE rather than phased using pairwise relationships.

Second, pedigree reconstruction within HPG studies provides trios and other informative pedigree structures that can be leveraged for many use-cases. We used the 2,602 reconstructed trios to find 3,415 DNMs and tracked known disease-causing mutations through extended pedigrees. Pedigrees and relationships are also particularly useful for tracking transmission of rare variants, providing increased confidence in variant calls and allowing for the use of more traditional Mendelian genetic analyses. Pedigrees can be particularly useful when combined with follow-up chart reviews and the ability to recontact patients and their family members.

We show that cryptic family structure in a large sequencing dataset presents an opportunity to harness a valuable, untapped source of genetic insights rather than a nuisance that must be managed during downstream analyses. As we enter the era of genomic-based precision medicine, we see a critical need for additional innovative methods and tools that are capable of effectively mining the familial structure and distant relatedness contained within the ever-growing sequencing cohorts.

### Description of Supplemental Data

Additional methods, thirteen tables, eight figures, and one excel file.

## Conflict of interest

The study was funded by Regeneron Pharmaceuticals. In addition to many of the authors being employed by and stockholders in Regeneron Pharmaceuticals, G.D.Y. is a cofounder and a member of the board of directors of Regeneron Pharmaceuticals. J.G.R., J.C.S., C.S. and L.H. are listed inventors on a related provisional patent application filed by Regeneron Pharmaceuticals (62/555,597), which discloses the simulation framework and methods for identifying the familial relationships, phasing the pCHMs, and calling the DNMs. Additional information for reproducing the results described in the article is available upon reasonable request and subject to a data use agreement.

### Acknowledgments

We thank the MyCode Community Health Initiative participants for their permission to use their health and genomics information in the DiscovEHR collaboration.

## Web resources

DeNovoCheck - https://sourceforge.net/projects/denovocheck

FBAT - https://www.hsph.harvard.edu/fbat/fbat.htm

GATK - https://software.broadinstitute.org/gatk/

QTDT - http://csg.sph.umich.edu/abecasis/qtdt/

SimProgeny - https://github.com/rgcgithub/SimProgeny

TopMed - https://www.nhlbiwgs.org

